# Balancing skill against difficulty - behavior, heart rate and heart rate variability of shelter dogs during two different introductions of an interactive game

**DOI:** 10.1101/838524

**Authors:** Christine Arhant, Bernadette Altrichter, Sandra Lehenbauer, Susanne Waiblinger, Claudia Schmied-Wagner, Jason Yee

## Abstract

Interactive games may boost positive well-being by combining the benefits of rewards with cognitive and social enrichment. While a gradual introduction to the game can promote greater learning and skill, a hasty introduction can lead to low success and frustration. Here, we examine two methods of introducing an interactive game to dogs (*Canis lupus familiaris*) to test whether they elicit differences in success rate, stress-related behavior, and autonomic regulation of the heart.

Twenty-eight dogs living in shelters were given the opportunity to play with an interactive game that consists of four boxes with different opening mechanisms. Dogs were introduced to the interactive game in one of two ways: gradually vs hastily. Gradual introduction consisted of allowing the dog to first play a partial (2 out of 4 boxes) version of the game with a human experimenter demonstrating the opening mechanism of the boxes twice, followed by exposure to the complete game. Hasty introduction consisted of the same procedures but presented in a different order, with the complete game presented before the partial version. Dog behavior was obtained via video recordings and pre- and post-game mean HR, RMSSD, SDNN, RMSSD/SDNN ratio were assessed using R-R intervals obtained with a Polar heart rate monitor (RS800CX). Linear mixed effects analyses (LMM) were calculated for success and behavior component scores and for change from pre- to post-game period in HR & HRV variables. In addition, HR and HRV parameters were analyzed with Pearson correlations.

Dogs introduced to the game in a gradual manner had a significantly higher rate of success compared to dogs introduced in a hasty manner (LMM: p < 0.001). Dogs introduced to the game gradually also displayed less stress related-behavior, e.g. displaying lower scores for the arousal (p < 0.001) and displacement (p < 0.001) components. Correlation analysis revealed a negative correlation between HR and RMSSD during baseline in all dogs (pre-game, day 1: gradual: r = −0.52; hasty: r = −0.72) that gradually transformed into a strong positive correlation in the gradual introduction group (post-game, day 2: r = 0.78), whereas it remained negative over all evaluation periods in the hasty introduction group (post-game, day 2: r = −0.83).

Overall, our findings on success rate, dog behavior, and HR/HRV suggest that the way a moderately difficult game is introduced plays a major role in determining how the experience of game play is perceived. Our results are consistent with the hypothesis that gradual introduction including demonstration promotes an enjoyable experience characterized by greater likelihood of reward, less stress-related behavior, and a physiological profile that may involve activation of both sympathetic and parasympathetic branches of the autonomic nervous system. We suggest that this may be a physiologic signature of successful achievement and that a learning experience in which skills are balanced against difficulty promote pleasant emotional states.

## 1. Introduction

Dogs benefit from an environment enriched with toys or feeding enrichment [1-3] as well as from interactions with humans [4-7]. So-called interactive games or toys are designed for the joint application of dog and human and therefore combine the benefits of human interaction, food and cognitive enrichment [e.g. 2, 5, 8]. The basic idea behind interactive games is to solve a task and thereby gain access to a treat under the close supervision of a human. To get dogs accustomed to the game, the level of difficulty is gradually raised and it is recommended that dog owners provide assistance and encouragement to increase the level of involvement by the dog [e.g. 9].

Social learning is learning through observation of others, whether it be humans or other dogs [10, 11]. Various studies have shown that demonstrations by humans improve a dog’s performance especially in moving and manipulating objects [e.g. 12, 13, 14]. Therefore, adding human demonstration to the introduction of an interactive game should allow a dog to solve the tasks quicker and increase the benefit of interactive game use as success in a task has been shown to be a catalyst for positive emotion [15]. In contrast, a hasty introduction without human demonstration might lead to frustration and distress [15, 16].

Measuring stress and assessing welfare, particularly the valence of emotional states in animals, is challenging. Most approaches rely on multiple parameters integrating behavior and physiology. The measurement of heart rate (HR) and heart rate variability (HRV) is an important non-invasive tool to assess the autonomic impact of different types of stimuli in animals [17]. HRV is the variation in time intervals between consecutive heartbeats and can be quantified by analyzing R-R or inter-beat intervals. Since it is influenced by sympathetic and parasympathetic neural activity, HRV provides an insight into the activity of the autonomic nervous system [18, 19] and has been used to assess emotional state of dogs confronted with different types of stimuli [20-22]. Increases in HR can be regarded as a measure of arousal, with either positive or negative emotional valence [23] and HR is a product of influences from both branches of the autonomic nervous system [24]. Increases in HRV, particularly the high frequency component and respiratory sinus arrhythmia, can be regarded as a measure of parasympathetic activity [19, 25]. Two commonly used HRV parameters are the time-domain measures SDNN (standard deviation of the R-R, or N-N, intervals), which is influenced by sympathetic and parasympathetic activity, and RMSSD (root mean square of successive differences) which estimates short-term components of high-frequency beat-to-beat variation and therefore reflects parasympathetic activity [17]. In dogs, pharmacological and surgical blockade showed that both the SDNN and an estimate of the high frequency component of the R-R interval variability (‘vagal tone index’) seem to be largely controlled by parasympathetic influences [24].

The aim of this study was to compare two different styles of introduction to an interactive game on the degree of success or reward-acquisition and the valence of emotional states experienced during game play. Whereas a gradual approach with a demonstration of opening mechanisms is recommended by manufacturers, inexperienced dog owners may use a hasty introduction lacking social demonstration. We hypothesize that dogs given a gradual introduction will be more successful in reaching the treats and will show fewer signs of stress-related behavior and lower arousal (as measured by HR).

## 2. Animals, Materials & Methods

In this study, shelter dogs were tested in two conditions (stepwise (s), complete (c)) in a counterbalanced order, mimicking a gradual (order s/c) and a hasty introduction (order c/s), with the objective to compare dog behavior, heart rate (HR) and heart rate variability (HRV). The interactive game “Poker Box 1 ©Trixie” containing four boxes with different opening mechanisms from the company TRIXIE Heimtierbedarf GmbH & Co. KG was used (Figure 1a). The stepwise condition represented a slow introduction into the game including a repetition of movements while filling the game to demonstrate the opening mechanisms of the boxes. The complete condition represented the opposite, a rather hasty approach, which did not involve any help through additional demonstration of opening mechanisms of the boxes. The study was discussed and approved by the institutional ethics and animal welfare committee in accordance with GSP guidelines and national legislation.

**Figure 1:**
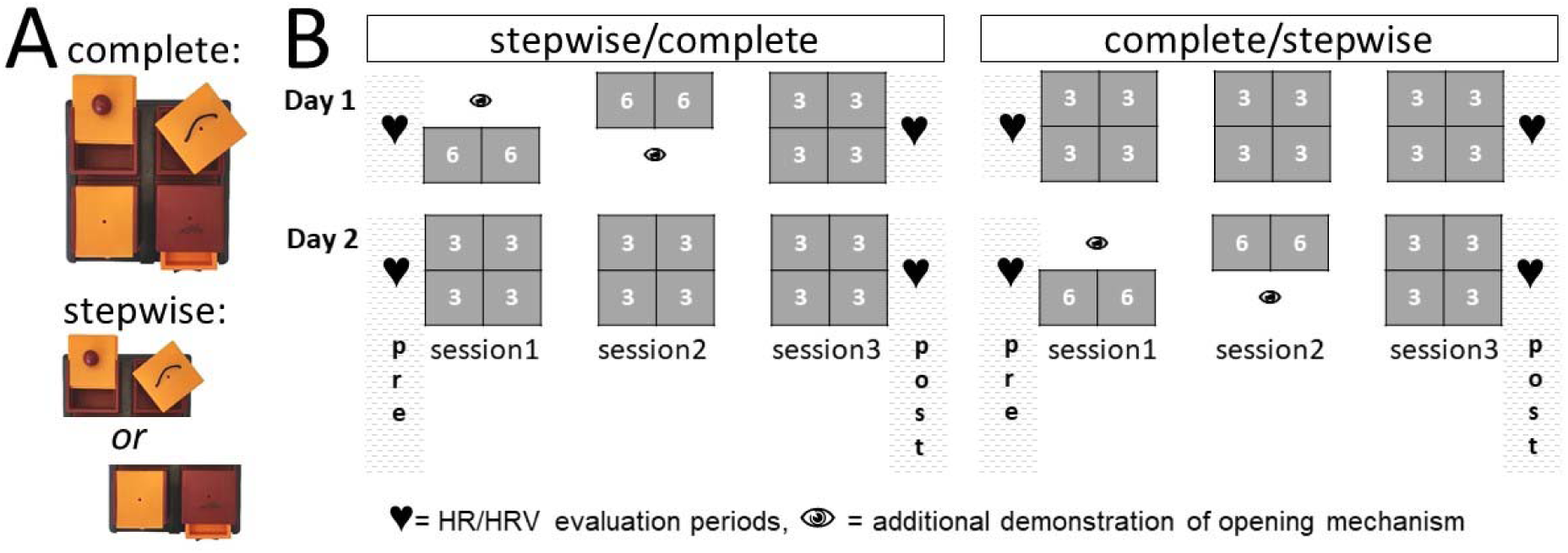
Interactive game used in the study (A) and layout of the experimental procedure (B)

### 2.1. Animals

28 dogs (17 females, 11 males; mean age ± SD: 4.5 ± 3.2 years) were tested in both conditions in a counterbalanced order. Twenty dogs were mixed breeds and eight were of a recognized breed (1 Rottweiler, 1 Pit Bull Terrier, 1 American Staffordshire Terrier, 1 Jack Russel Terrier, 1 Akita Inu, 1 American Bulldog, 2 American Staffordshire Terrier). For the duration of the study all dogs were housed in a local animal shelter and handled by the two experimenters (BA & SL) and their usual caretakers.

### 2.2. Experimental procedure

Each condition included three consecutive sessions of ‘playing’ an interactive game of a maximum duration of ten minutes with two breaks of five minutes between sessions (Figure 1b). Session 1 and session 2 of the stepwise condition differed from session 1 and session 2 of the complete condition. The setup of session 3 was identical in both conditions. The dogs were randomly assigned to one of two orders of conditions (stepwise/complete (s/c), complete/stepwise (c/s)). Dogs within order s/c (15 dogs) started with the stepwise condition (mimicking a gradual introduction as recommended by the manufacturer) while dogs within order c/s (13 dogs) started with the complete condition (simulating a hasty approach by the owner). The two conditions were carried out on two separate days with usually one resting day between conditions (Figure 1b).

In the stepwise condition, in the first session the dog was introduced to only two out of the four different boxes of the interactive game (Figure 1a – stepwise). Six trials with each box filled with one treat per trial were conducted (Figure 1b). In total, 12 treats could be obtained. While filling the boxes, the experimenter (SL) first called the dog’s name and showed the treat to the dog. In addition during the stepwise condition, the experimenter demonstrated the opening and closing of the boxes twice. After a five-minute break the next session started. In session 2 the setup was identical to session 1 but the other two, still unknown boxes were presented to the dog. The two boxes were always presented simultaneously. In the third session all four boxes were used simultaneously without an additional demonstration of opening mechanisms. Three trials with one treat per box were carried out (12 treats could be obtained). In the complete condition all sessions (1, 2 and 3) had the same setup as the third session in the stepwise condition. While the boxes were filled, the dog, accompanied by the handler (BA), watched from the waiting area (a blanket) opposite from the game until the experimenter gave the start signal.

The experiment was carried out in a wooden detached house surrounded by a fenced garden located on the grounds of the animal shelter. The testing room was equipped with a blanket that served as the waiting area for dog and handler during the sessions, a water bowl, an additional soft resting place for the dog and a couch. The interactive game was presented opposite to the waiting area on a non-slip mat. During the entire experiment, video recordings of dog behavior were conducted (four digital cameras placed in the corners of the room: GV-BX 1300; software: GEO Vision Digital Surveillance System 8.5.9.09). Inter-beat intervals (IBIs) were recorded using a Polar^®^ heart rate monitor (RS800CX, Polar^®^ Electro Oy) previously validated for use in dogs [26, 27].

On testing days, the caretaker and the dog met with the experimenters for a short walk. After the walk and the fitting of the Polar^®^ heart rate monitor the caretaker left the dog with the experimenters. The transmitter of the Polar^®^ heart rate monitor was placed ventrally on the chest area with the electrodes extending laterally to each side [26]. The dog was allowed 10 minutes to habituate to the new environment. Two six-minute HR/HRV evaluation periods per testing day, one after habituation before the start of the first session (pre day 1/pre day 2) and one after the third session (post day 1/post day 2), were scheduled. During these periods, the dogs were kept on a short leash in a standing position.

### 2.3. Data handling

#### 2.3.1. BEHAVIOR

The software Solomon Coder (Copyright © 2016 by András Péter) was used for a detailed analysis of behavioral elements of dogs during the test sessions. Continuous sampling was used to record frequencies and durations of dog behavior. Coding was carried out by one trained observer (handler, BA). For a description of all coded behaviors see the ethogram (Table 1). The duration of a session depended on how quickly the dog succeeded to open the boxes. A session started with the *start signal* (the start signal in both conditions was nodding combined with saying ‘Such!’, the German word for ‘search’) of the experimenter and ended when the dog had accomplished all tasks of each trial or the *stop signal* was given by the experimenter. Each achieved treat was coded as ‘success’. The stop signal was given when the dog stopped interacting with the game and did not resume playing after a previously determined method of encouraging: in the stepwise condition the experimenter pointed at the game saying the word ‘Such!’ repeating this a maximum of six times every ten seconds. The procedure was the same in the complete condition but without pointing at the game. For further analyses of behavior, we calculated a rate per minute for frequencies and a percentage of time for durations.

**Table 1:** Ethogram: Behavioural elements recorded during the interactive game sessions. Durations (dur) or frequencies (fr) were recorded.

| Behavioural elements | Description |
| --- | --- |
| **Interaction with the game** |  |
| Mouth (dur) | Dog is manipulating the game with the mouth and/or teeth |
| Paw (dur) | Dog is manipulating the game with front legs |
| Mouth & Paw (dur) | Dog is manipulating the game with the mouth/teeth and front legs simultaneously |
| **Contact with human** |  |
| Look at experimenter (dur) | Dog is looking at the experimenter |
| Look at handler (dur) | Dog is looking at the handler |
| Body contact experimenter (dur) | Dog is sitting, lying or moving with body contact to the experimenter |
| Body contact handler (dur) | Dog is sitting, lying or moving with body contact to the handler |
| **Behaviour during play** |  |
| Look away, call (dur) | Dog is averting sight or turning head away when called by the experimenter |
| Scratching (dur) | Scratching the own body with one of the two hind legs |
| Lip lick (fr) | Part of tongue is shown, short movements along the upper lips |
| Shaking (fr) | Rotation movement of the whole body |
| Stretching (fr) | Different body parts (front legs, hind legs, back and neck) in stretched posture |
| Play bow (fr) | The front part of the dog's body is lowered - lying on the front legs and the posterior is heightened - dog is standing on hind legs |
| Yawning (fr) | Mouth opened widely, the tongue is rolled up, sometimes accompanied by a high, soft noise |
| Barking (dur) | Short, low frequency noises |
| Whimpering (dur) | Soft, high pitched vocalisation |
| Tail wagging (dur) | Repetitive movements of the tail |
| Panting (dur) | Mouth is opened, increased frequency of in- and exhaling, often with a hearable breathing noise |
| **Location** |  |
| Around mat (dur) | Dog is standing, walking or sitting on the rubber mat, where the game is placed upon |
| At door (dur) | Dog is sitting or standing close to the door of the testing room |

To determine intra- and interrater-reliability, different sessions of twelve dogs were coded twice. Videos were chosen randomly from the whole period of data acquisition. The intra- and inter-rater-reliability for all behavioral variables was determined by using the Spearman rank correlation coefficient. Intra-rater reliability was high (mean over all coded behaviors: r_s_ = 0.99, p < 0.01; minimum over all coded behaviors: r_s_ = 0.91, p < 0.01). Also the inter-rater reliability resulted in excellent agreement (mean: r_s_= 0.99, p < 0. 01; minimum: r_s_=0.85, p < 0.01).

#### 2.3.2. HR AND HRV

Polar^®^ Precision Performance was used for a preliminary assessment of the measurement periods’ error rates. First a rough estimation over the entire six-minute evaluation period was done. Two dogs with extremely high error rates (> 30 %, Polar^®^ filter power “very low”) were excluded from further analyses at this point (c/s = 2 dogs). In the next step, error rates were determined separately for every minute and three consecutive minutes were chosen for manual error correction. If possible, the first three minutes were used. Only if other minutes with considerably lower error rates were available, these were selected. During the evaluation of error rates per minute, three dogs had to be excluded from further analyses as it was not possible to find three consecutive minutes with acceptable error rates (s/c = 1, c/s = 2). In the third step, the time interval and IBI data of the selected minutes were exported to an Excel file and errors were identified visually or based on differences between consecutive IBIs and then corrected manually based on a commonly used approach (Marchant-Forde et al. 2004, Gamelin et al. 2008, Jonckheer-Sheehy et al. 2012, Schöberl et al. 2014, Giles et al. 2016, Lensen et al. 2017). Details on error correction are provided in the supplementary material. Errors in data of 23 dogs were corrected manually and only three-minute recordings with less than 10 % errors were included in the statistical analyses. Further three dogs had to be excluded (s/c = 2, c/s =1) and further three dogs did not fulfill this criterion for all evaluation periods (s/c = 2, c/s =1). The final sample for HR/HRV analyses with data in all evaluation periods consisted of 17 dogs (s/c = 10, c/s = 7); data for post day1/post day 2 was available in 20 dogs (s/c = 12, c/s = 8). The corrected three-minute recordings of IBI data were imported in the software Kubios HRV Standard Version 3.0.2 (Kubios Oy). The following time domain parameters (mean HR, RMSSD, SDNN, RMSSD/SDNN ratio) were used for statistical analyses.

### 2.4. Statistical analyses

Descriptive and inferential statistical analyses were carried out with the statistical package SPSS Statistics Version 22.0 and 25.0 (IBM Corp 2013) except for linear mixed effects analyses (LMM) which was performed using the statistical software R (R Core Team 2018) and “lme” function from the package “nlme” [28].

In order to group dog behaviors during play with the strategy game, principal component analyses (PCA) followed by Varimax rotation were carried out. Bartlett’s test of sphericity was required to be significant and the Kaiser-Meyer-Olkin criterion should be at least 0.5. Components were required to have an eigenvalue greater than one. To include behaviors in the final solution, the Anti-Image Correlation Matrix diagonal was required to be at least 0.5 and behaviors with double loadings larger than 0.4 were excluded from further analyses. Initially, all dog behaviors during game use including contact with humans were analyzed in one PCA. However, interactions with humans resulted in several double loadings higher than 0.4. Therefore, interactions with humans were analyzed separately. Only one variable (barking) did not fulfil our conditions and was excluded from the final solution. Factor scores were calculated by using the Regression scores method of SPSS. The PCA of dog behavior during game use resulted in a four components solution explaining 62.3% of the total variance (see table 2 for factor loadings; % variance explained by factor score fiddle: 20.2; arousal: 17.5; avoidance: 13.4; displacement: 11.1). The second PCA regarding contact with humans resulted in a one component solution explaining 52.6% of the total variance (see table 2 for factor loadings).

**Table 2:** Factor loadings and Spearman Correlations between success during play with an interactive game and dog behavior based on data from 28 dogs tested in six sessions over two conditions (in total: 168 sessions).

| | Factor loading | Success/<br>minute<br>$r_s$ | p |
| --- | --- | --- | --- |
| **Interactions with game** |  |  |  |
| % mouth | -- | 0.88 | < 0.001 |
| % paw | -- | 0.46 | < 0.001 |
| % mouth & paw | -- | 0.61 | < 0.001 |
| % time interacting with game | -- | 0.91 | < 0.001 |
| % around mat | -- | 0.74 | < 0.001 |
| % at door | -- | -0.42 | < 0.001 |
| **Dog behaviour** |  |  |  |
| *Factor score fiddle* |  | 0.26 | 0.001 |
| Stretch/min | **0.935** | -0.10 | 0.204 |
| Playbow/min | **0.932** | -0.11 | 0.176 |
| *Factor score arousal* |  | -0.49 | < 0.001 |
| % whimpering | **0.804** | -0.27 | < 0.001 |
| Liplick/min | **0.717** | -0.61 | < 0.001 |
| % tail wagging | **0.597** | -0.34 | < 0.001 |
| *Factor score avoidance* |  | -0.58 | < 0.001 |
| % lookaway when called | **0.761** | -0.48 | < 0.001 |
| Yawning/min | **0.691** | -0.27 | < 0.001 |
| % panting | **0.581** | -0.44 | < 0.001 |
| *Factor score displacement* |  | -0.34 | < 0.001 |
| Shaking/min | **0.808** | -0.32 | < 0.001 |
| % scratching | **0.764** | -0.15 | 0.054 |
| Not included in final solution: |  |  |  |
| % barking |  | -0.22 | 0.004 |
| **Behaviour towards humans** |  |  |  |
| *Factor score contact humans* |  | -0.88 | < 0.001 |
| % looking at experimenter | **0.767** | -0.83 | < 0.001 |
| % looking at handler | **0.854** | -0.82 | < 0.001 |
| % bodycontact to experimenter | **0.641** | -0.73 | < 0.001 |
| % bodycontact to handler | **0.611** | -0.54 | < 0.001 |

To check whether LMM fulfilled the model assumptions, residual plots of all linear mixed effects models were obtained and inspected graphically for normality and homogeneity of variances. When outliers were identified (> 3 SD), the analyses were repeated without them and if there were no obvious differences in the results, the models based on the complete dataset are presented (which was the case for all models). If model assumptions were not fulfilled (e.g. variance inhomogeneity) the dependent variable was transformed using a log or rank transformation. This was necessary for some behavior factor scores. The transformation which led to the model that fulfilled the assumption of variance homogeneity best was chosen (‘fiddle’: log; ‘avoidance’: rank; ‘displacement’: log).

Regarding behavior all linear mixed models were calculated with order (s/c, c/s) and condition (stepwise, complete) as fixed factors and order*condition as two-way interaction. As random effect the identity of dogs was included. As several models were calculated (‘success’, five behavior factor scores) a correction for multiple testing using the Bonferroni method was done in the cluster of 5 tests including all behavior factor scores: the p - value considered to be significant was p < 0.01.

To analyze HR and HRV, the differences (deltas Δ) between pre- and post-test evaluation periods were calculated. The deltas were obtained by subtracting pre from post-periods (post – pre on day 1 and day 2).Therefore a positive delta indicates an increase and a negative delta indicates a decrease in values after the third session. The LMM included the same fixed effects and random effect as the models for behavior.

In addition, HR and HRV parameters were analyzed with Pearson correlations for each order*condition combination separately. To detect possible differences between dogs in the order s/c or c/s at the start of testing, pre values were compared between dogs on day 1 and day 2 by using t-tests.

## 3. Results

### 3.1. Descriptive statistics of duration of use and interaction with the game

Overall, a session including time waiting for the game to be filled with treats and time ‘playing’ with the game had a mean duration of 4.4 ± 1.9 min (Minimum: 1.4 min; Maximum: 9.5 min). For dogs in the recommended order s/c the time manipulating the game in the stepwise condition on the first day was on average 2.6 ± 1.2 min per session and in the complete condition on the second day 1.9 ± 0.9 min per session. Dogs in the order c/s engaged shorter with the interactive game (complete: 1.0 ± 1.0 min, stepwise: 1.7 ± 1.1 min per session). Overall, dogs manipulated the game mainly with the mouth. The proportion of time staying around the mat differed slightly between the two orders and dogs in order c/s stayed around the mat less and stood at the exit door of the experimental room longer in the complete condition (see Table 3). The dogs in order s/c reached on average 10 ± 4 treats per session on the first day and 11 ± 4 treats per session on the second day (in session 3 all dogs obtained 12 treats except one that did not perform at all). Dogs in order c/s were less successful and gained on average 2 ± 3 treats per session on the first day and 8 ± 5 treats per session on the second day. Sessions were interrupted more frequently in the order c/s (day 1: 3 ± 1, day 2: 2 ± 2; order s/c – day 1: 1 ± 2, day 2: 0 ± 1).

**Table 3:** Percentage of time spent interacting with the game by order and condition

|  | s/c |  |  |  | c/s |  |  |  |
| --- | --- | --- | --- | --- | --- | --- | --- | --- |
|  | stepwise |  | complete |  | complete |  | stepwise |  |
|  | Mean | SD | Mean | SD | Mean | SD | Mean | SD |
| % mouth | 51.8 | 18.6 | 66.3 | 24.9 | 23.2 | 17.1 | 47.1 | 29.8 |
| % paw | 6.1 | 6.5 | 8.1 | 9.6 | 0.4 | 1.2 | 3.0 | 6.7 |
| % mouth & paw | 4.1 | 4.3 | 5.0 | 4.9 | 1.4 | 4.9 | 3.5 | 5.5 |
| % time interacting with game | 62.0 | 22.2 | 79.4 | 25.5 | 25.1 | 21.1 | 53.5 | 33.7 |
| % around mat | 92.2 | 9.2 | 94.5 | 9.8 | 80.6 | 14.1 | 86.7 | 18.2 |
| % at door | 0.6 | 1.8 | 0.6 | 2.3 | 1.8 | 3.1 | 0.4 | 1.2 |

### 3.2. Effect of game setup on rate of success

The linear mixed model analyses resulted in significant main effects for order and condition as well as a significant interaction of order*condition on the rate of success (Table 4). In particular, the success rate was much higher for dogs tested within order s/c as compared to dogs tested within order c/s on both the first and second test day (Figure 2 A).

**Table 4:**
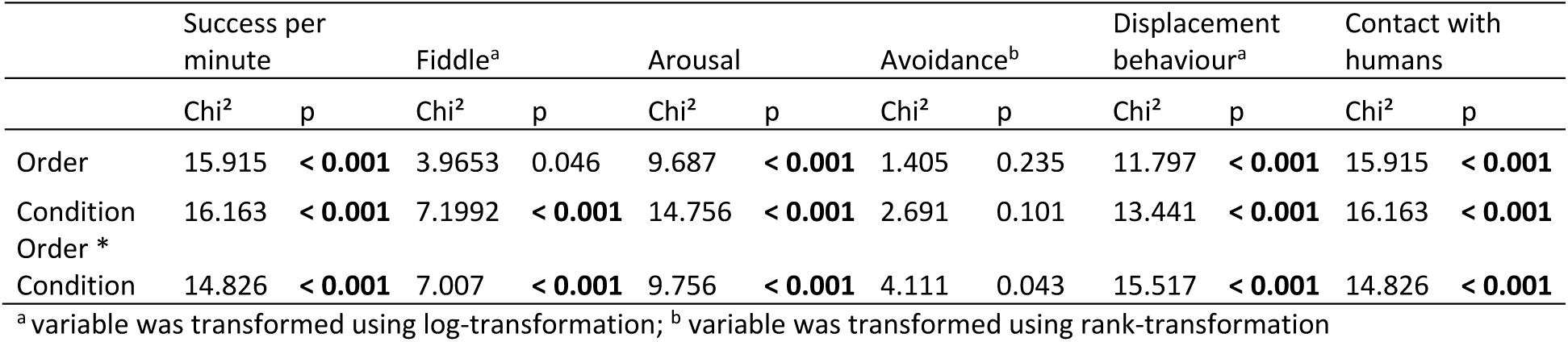
Results of LMM for effect of game set up on success rate and behavior factor scores. P-values remaining significant after multiple testing are in bold print.

**Figure 2:**
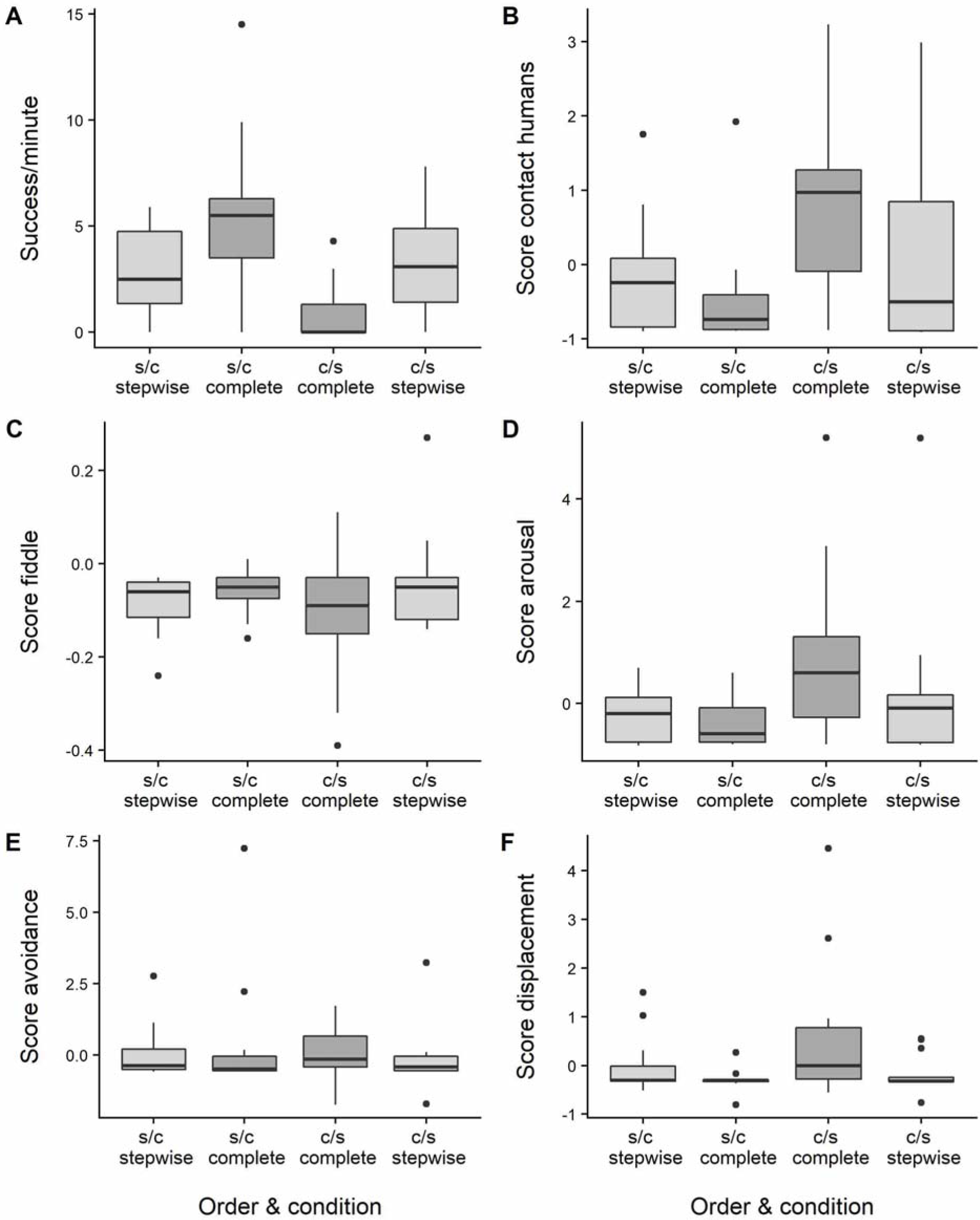
Boxplots of success per minute and behavior factor scores by order and condition

### 3.3. Effect of game setup on dog behavior

Significant main effects of order and condition and their two-way interaction on the factor scores of dog behavior (arousal, displacement) and contact with humans were found (see Table 4). Factor scores for arousal, displacement and contact with humans show a similar pattern (Figure 2B, D, F). Especially on day 1, dogs tested within order c/s displayed more behaviors indicative of high arousal, made contact more often with humans and displayed more displacement behaviors in contrast to dogs tested within the order s/c. These behaviors were generally observed more often on day one than on day two, but dogs tested in the order s/c displayed overall lower levels. For the factor score ‘fiddle’ only the main effect of condition and the interaction order * condition remained significant after correction for multiple testing (Table 4). The pattern observed contrasts the results regarding arousal, displacement and contact with humans (Figure 2 C). Behaviors included in the fiddle factor score were observed more often on day two of testing and the change in behavior was more pronounced in dogs within order c/s. No significant effects on the avoidance factor score were found (Table 4, Figure 2E).

### 3.4. Relationship of rate of success with dog behavior and use of interactive game

Increased success per minute was highly related to an increased proportion of time interacting with the game as well as manipulating the game with the mouth (see Table 2). The most prominent relationships between success and dog behavior was found for interactions with humans. The more dogs looked at or initiated body contact to the experimenter or handler the less successful they were. Regarding the other dog behaviors, lip-licking and avoiding interaction with the game, i.e. looking away when being called by the experimenter to resume playing and a high arousal factor score resulted in the highest negative relationship with success (Table 2).

### 3.5. Effect of game setup on HR and HRV

To ascertain that dogs in the two orders did not differ in absolute values of HR and HRV parameters, we compared pre-testing values (before interactive game use) on day1 (HR: s/c = 115±9, c/s = 123±18; SDNN: s/c = 55±15, c/s = 57±20; RMSSD: s/c = 44±18, c/s = 52±26) and day 2 (HR: s/c = 118±11, c/s = 125±22; SDNN: s/c = 57±8, c/s = 60±31; RMSSD: s/c = 56±15, c/s = 57±36) and found no significant differences between s/c dogs and c/s dogs (p > 0.3 for all tests).

The LMM resulted in no significant effects for any of the main effects (order (o), condition (c)) or the two-way interaction (order*condition (o*c)) for the delta values of HR (o: p = 0.693, c: p = 0.549, o*c: p = 0.755; s/c – s: −3.63 ± 9.38, s/c – c: −2.92 ± 5.80, c/s – c: −4.36 ± 8.97, c/s – s: −6.57 ± 6.83), SDNN(o: p = 0.999, c: p = 0.869, o*c: p = 0.575; s/c – s: 4.70 ± 22.33, s/c – c: −0.70 ± 15.13, c/s – c: − 0.71 ± 21.62, c/s – s: −2.29 ± 15.53) and RMSSD (o: p = 0.842, c: p = 0.579, o*c: p = 0.253; s/c – s: 12.26 ± 28.15, s/c – c: −0.26 ± 24.93, c/s – c: 2.20 ± 27.47, c/s – s: −5.23 ± 26.19). The LMM for the delta value of the RMSSD/SDNN ration resulted in a statistical tendency for the interaction order*condition (Chi^2^ = 2.75, p = 0.097; s/c – s: 0.14 ± 0.21, s/c – c: 0.00 ± 0.22, c/s – c: −0.01 ± 0.10, c/s – s: −0.08 ± 0.19); main effects were not significant (o: p = 0.912, c: p = 0.423). However, when correlations between HR and the two HRV parameters were analyzed separately for each evaluation period and both orders of testing, a substantially different pattern of correlation depending on the order of testing emerged (Figure 3 and 4). In the order c/s, the baseline measurement (pre day 1 (N = 7)) showed a negative correlation between HR and RMSSD (r = −0.719, p = 0.07) or HR and SDNN (r = − 0.719, p = 0.07). This negative correlation got stronger and statistically significant (p < 0.05) and remained stable over the further three time-points of HR and HRV measurements (post day 1 (N = 8): RMSSD r = −0.890, SDNN r = −0.879; pre day 2 (N = 7): RMSSD r = −0.813, SDNN r = −0.844; post day2 (N = 8): RMSSD r = −0.831, SDNN r = −0.818). In the order s/c, the correlations of the baseline measurements (pre day 1 (N = 10)) were similarly negative (HR-RMSSD r = −0.520, p = 0.12, HR-SDNN r = −0.601, p = 0.07). However, in the order s/c, this negative correlation got weaker over time (post day1 (N= 12): RMSSD r = −0.411, SDNN r = −0.434; pre day 2 (N = 10): RMSSD r = 0.081, SDNN r = −0.152), and finally turned into a statistically significant positive correlation on day 2 after playing the game (post day2 (N = 12): RMSSD r = 0.784, SDNN r = 0.707, p < 0.05). This pattern in order s/c was still present (Figure 3), when only dogs with data on all time points were included and the only dog in this order that did not successfully use the interactive game was excluded (N = 9, pre day 1: RMSSD r = −0.554, SDNN r = −0.631; post day 1: RMSSD r = −0.513, SDNN r = −0.484; pre day 2: RMSSD r = −0.170, SDNN r = −0.439 – all p-values > 0.05; post day2: RMSSD r = 0.685, p = 0.042, SDNN r = 0.485, p = 0.166). The correlation between SDNN and RMSSD was always positive and above r = 0.9 in both orders except for the reduced dataset in order s/c at the evaluation period post day 2, where it was slightly below (r = 0.884, p = 0.002, N = 9). Scatterplots of the baseline values of HR and RMSSD (pre day 1) and the measurement after interacting with the game on the second day (post day 2) show the development of HR and RMSSD for individual dogs over time (Figure 4).

**Figure 3:**
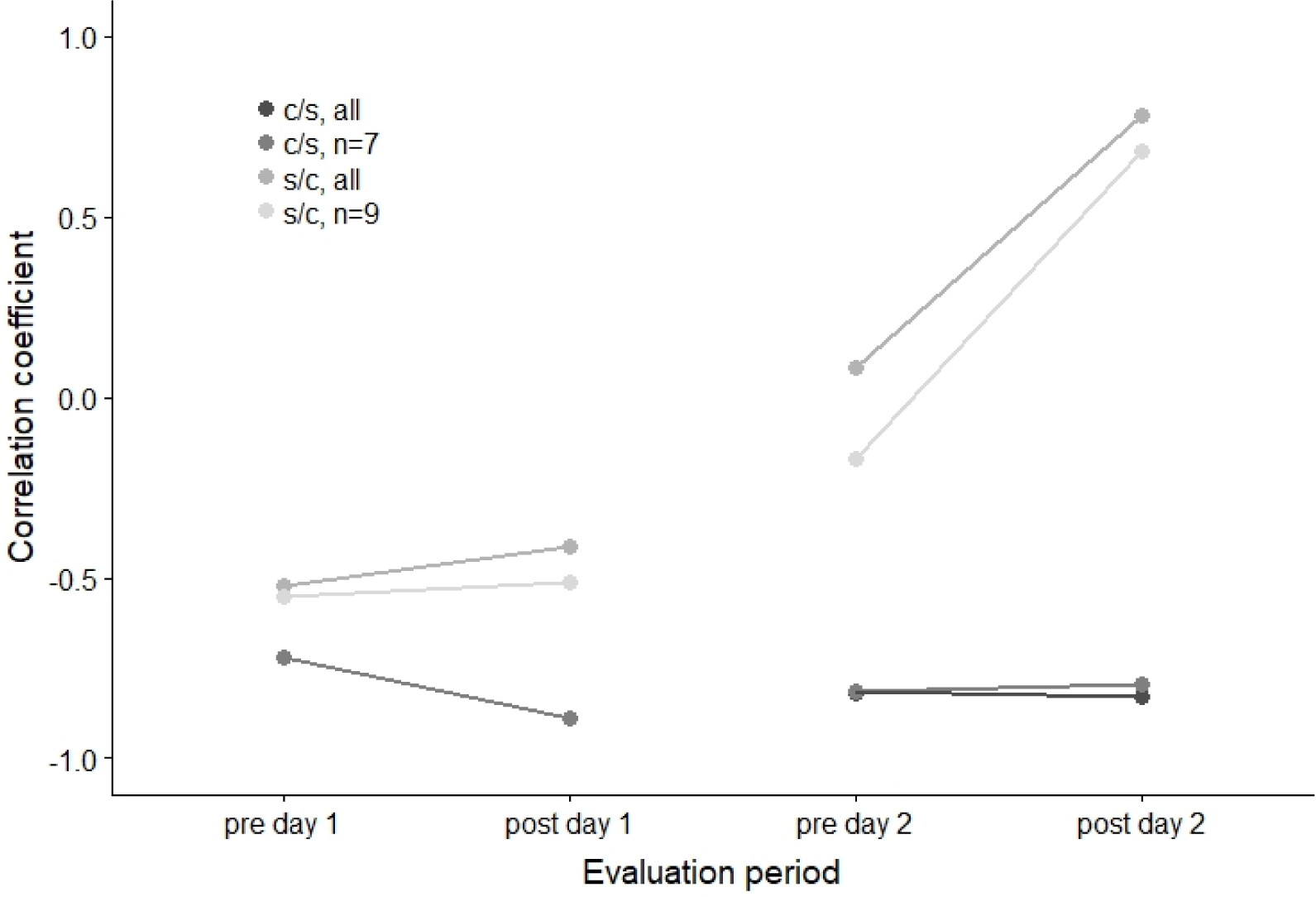
Development of Pearson correlation between heart rate (HR) and the heart rate variability parameter RMSSD over time by order (s/c & c/s). Due to missing values in the pre-evaluation periods, we display both the correlation coefficients based on the whole dataset (all) and the samples with complete data in all four evaluation periods.

**Figure 4:**
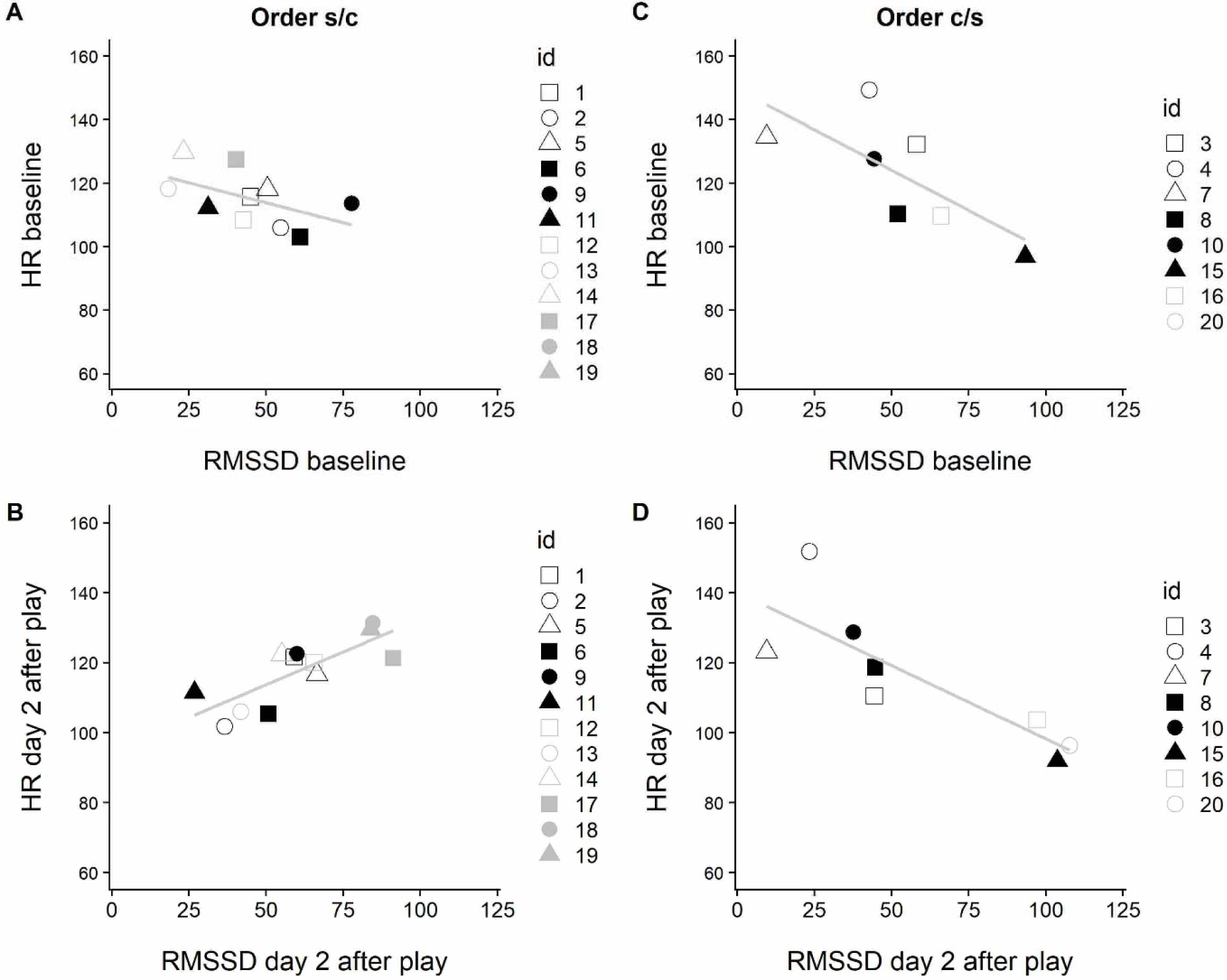
Scatterplots of the baseline values of HR and RMSSD (pre day 1) and after interacting with the game on the second day (post day 2) for individual dogs per order based on the whole data set

## 4. Discussion

In this study we investigated two different approaches to introduce an interactive game for dogs. A gradual presentation of the interactive game with human demonstrations of the opening mechanisms was compared to a hasty introduction without additional demonstrations of the opening mechanisms. We expected that the two introductions which differed in terms of difficulty and provision of human aid would lead to differences in success rate, occurrence of so-called stress-related behaviors and HR/HRV parameters. Regarding success and dog behavior, our results confirmed our hypotheses. Dogs given a gradual introduction had a significantly higher rate of success compared to dogs given a hasty introduction. Furthermore, for example the factor score ‘arousal’, which included the behaviors ‘whimpering’, ‘lip licking’ and ‘tail wagging’, decreased over time within both orders and followed a contrasting pattern compared to the rate of success. Whimpering or whining can be observed for example during separation from the owner, e.g. during kenneling [29-31] as well as during greeting of the owner [32] but also when dogs anticipate electric shocks [33]. Lip licking can frequently be observed as a response to a friendly but also a threatening approach by humans [34] or during physical contact with a human [21] but it was also found to be related to increased cortisol in hospitalized dogs [35]. Tail wagging is observed in a wide range of situations and towards positive and negative stimuli [36]. During operant learning tasks several different tail positions and movements were found to be related to success namely a high tail position, a non-wagging tail but also (short and quick) tail wagging [15, 37]. The three behaviors included in the arousal factor score are displayed in situations where the dog is aroused without necessarily indicating valence. In our case, we suggest that a low arousal factor score might indicate that the dogs were better able to focus on the task. Consequently, this led to a higher rate of success. The stepwise condition consisted of a gradual introduction to the task (only two of the four boxes at a time), including additional human demonstrations of the different opening mechanisms and use of communicative gestures such as pointing at the game. This set of differences compared to the hasty introduction clearly enhanced the dogs’ performance. It has already been shown that human demonstration and use of communicative gestures increase a dog’s performance in different tasks [13, 14, 38]. In our study setup, we were not able to tell which aspects contributed most to the increased performance of dogs. However, from an applied point of view, shelter dogs were more successfully engaged in this joint human-dog activity when introduced gradually to the game.

In particular in the early stages, interacting with shelter dogs can be challenging for staff, volunteers or researchers. In our study, both experimenters were unfamiliar to the dogs on day 1 of testing. On one hand, past studies reported that familiarity of the demonstrator did not affect success in a task [39]. On the other hand, increased attention during a problem-solving task only occurred towards familiar humans with whom the dog had a close relationship [40]. It has been shown that shelter dogs form attachment bonds with humans quickly [41] On day 2 of testing, a bond between the dogs and the handler/experimenter might have started to develop, enhancing the dog’s feeling of security and thereby improving it’s performance. In both orders dogs showed more behaviors indicative of high arousal or stress on day 1 with overall higher levels in the hasty approach with factor scores ‘contact with humans’, ‘displacement’ and ‘arousal’ following a similar pattern. Increasing experience with manipulating the game (and more success) was associated with lower levels of interactions with the humans present in both orders on the second day. When confronted with an unsolvable task dogs may give up and gaze at the human experimenter [e.g. 42, 43]. Initiating contact to humans in response to a potentially stress-inducing situation could be interpreted either as an attempt to get help from a human or represent a displacement or redirected behavior [44]. We conclude that the second testing day was less challenging for dogs in both orders but overall dogs given a gradual introduction were more successful and less aroused, stressed or frustrated. However, to clearly separate the effect of the two different conditions and orders and the two testing days, complementing the study design with two more orders of the conditions (stepwise/stepwise and complete/complete) would have been necessary.

Behaviors contained in the factor score “fiddle” reached higher levels on the second day in both orders. This factor score contained two behaviors: the play bow and stretching. The play bow indicates intention to play [45] and possibly reflects positive emotions [46]; stretching may indicate relaxation [47] but has also been associated with restricted housing conditions and single housing [48, 49]. When compared with our other findings, “fiddle” factor scores follow the pattern of the rate of success and contrast the pattern of the factor scores “arousal”, “displacement” and “contact to humans”. This may indicate that in our study, the factor score “fiddle” represents behaviors related to an underlying emotional state of positive valence. This is supported by findings that successful problem-solving can cause positive emotions [15, 50]. In conclusion, the results of behavior analyses show that the gradual approach led to less behaviors likely to indicate negative emotions and to more behaviors likely to indicate positive emotions in this context.

We expected dogs’ physiology to reflect, whether as a cause or a consequence, differences in behavior elicited by the way in which the interactive game was presented. Our initial hypothesis was that dogs would exhibit greater HR and lower HRV when given a hasty introduction, consistent with the notion that heightened stress from the frustration of unsuccessful usage of the game would lead to heightened arousal. While this notion was supported by behavioral observations (behavioral arousal was negatively correlated with success rate), neither condition nor order produced significant differences in the change of HR, SDNN, or RMSSD from before to after play, suggesting dogs experienced equivalent levels of physiological arousal regardless of how the interactive game was presented. This mismatch between behavioral and physiological arousal may be due to underestimating the physiological arousal that can be associated with successful play attempts. According to the two-dimensional model of core affect [51, 52], arousal and valence are orthogonal dimensions. Therefore, the order of interactive game presentation, even if it produced divergent experiences of stress or enjoyment, could both be equivalently arousing. Further, while behavior was observed directly during interacting with the game, the physiological data were taken somewhat later. Potential differences may have declined already. This finding also highlights the need to take caution when drawing conclusions on arousal based solely on physiological measurements. While order of presentation did not influence physiological arousal as measured by change in heart rate and heart rate variability, our findings on success rate and stress-related behavior suggested that order played a major role in determining the valence of the experience (i.e. whether stressful or enjoyable). In addition, our behavioral findings are supported by LMM resulting in a statistical tendency for the interaction between the order of testing and the condition of game play regarding change of RMSSD/SDNN ratio. The lack of findings regarding the effect of the game setup on delta values of the two single HRV parameters RMSSD and SDNN could on the one hand indicate that effects were too small to be shown with this low sample size (sample size was reduced from 28 to 17 because of high errors in R-R measurements), on the other hand looking at the change of single parameters might not tell the whole story [53, 54]. Thus, we performed secondary analyses and found a substantial change in the pattern of correlation between HR and the HRV. In general, HR is negatively correlated to HRV for physiological and mathematical reasons [24, 55]. As heart rate increases the interval between successive beats shrinks, resulting in less potential for variation between inter-beat intervals, and thus lower HRV. For this negative correlation between heart rate and heart variability to be overcome (and transform to a positive correlation) requires a shift in the central regulation of cardiac autonomic activity, one in which parasympathetic activity is elevated against a background of heightened HR. This change of pattern suggests a pattern of dual autonomic activation, or concurrent activity of the sympathetic and the parasympathetic branches of the autonomic nervous system [56]. The results of correlating HR and HRV demonstrate two important things. First, in all dogs there is a negative correlation between HR and HRV during baseline (i.e. pre day 1), consistent with what is known about cardiac autonomic regulation during affectively neutral or negative contexts [24, 55]. Second, during the last measurement after game use on day 2, several dogs that were first presented with the game in a stepwise fashion with human assistance experienced both moderate to high HR with increased HRV, in particular increased RMSSD, while dogs that were first presented with the complete game without human assistance persisted in displaying a negative HR:HRV correlation. Moreover, as the dogs first given a stepwise introduction with human assistance progressed through the experiment, we observed a gradual shift in HR:HRV correlation, from negative to positive. Our finding bears similarity to a study on cognitive enrichment in pigs, which initially led to increases in HR but not RMSSD. After two weeks pigs had to press a button five times instead of once to receive the food reward and the increase in skill and effort was associated with increases in RMSSD [57]. We suggest that concurrent activation of the sympathetic and parasympathetic branches of the autonomic nervous system might be a physiological signature of successful achievement in animals.

In humans, a pleasant state of absorption during a challenging activity, with an optimal and escalating balance between the skills of the person and the demands of the activity, is described as “flow” [58, 59]. Based on the results regarding success in obtaining treats (all 12 treats were obtained in session 3 on day 2 in the s/c group), our gradual introduction may have achieved a good balance between the dogs’ skill and the demands of the task. Studies investigating the flow-experience in humans during a stressful event found the highest self-reported flow values with moderate activation of the sympathetic branch (low frequency HRV) and moderate to high activation of the parasympathetic branch (high frequency HRV) [60, 61]. The relationship between activation of the sympathetic nervous system and the flow-experience was described as an inverted u-shape, i.e. very low and very high activation of the sympathetic branch reduced the “flow-experience” [60]. We hypothesize that our stepwise introduction balanced skills of the dogs against the demands of the task, and may have induced a ‘flow-like state’ in some dogs. While generalization of human constructs like flow to non-human animals must be handled with extreme caution, iterative escalation between skill and challenge to achieve success on a task is likely to be a basic phenomenon not limited to humans. To the extent that they are capable of achieving mastery at a task that aids their survival, the study of flow-like states in non-human animals promises to offer novel insights into how to provide animals with more optimal experiences and improved welfare.

Overall, we suggest that using interactive dog games in a way so that the demands of the task are balanced against the skills of the dog, neither resulting in boredom nor excessive demands but representing ‘an optimal challenge’, has the potential to induce a pleasant ‘flow-like’ emotional state in animals. Further studies are needed to verify whether ‘flow-like’ states exist in animals.

## 5. Conclusion

A gradual introduction to an interactive game combined with a demonstration of the possible mechanisms to solve the tasks led to a higher rate of success and a decreased occurrence of stress-related behaviors during play in a new environment with unfamiliar humans. Although, neither condition nor order did have influences on shelter dogs’ heart rate and heart rate variability change from before to after using the interactive game, the patterns of correlation of HR and HRV in the group with a gradual introduction to the game further suggest that some dogs experienced a pleasant emotional state during/after the use of the game. The stepwise approach not only facilitated success and limited behavioral signs of arousal, stress or frustration but very likely induced positive emotions possibly similar to the flow-experience in humans. Therefore to maximize the benefits, it is of high importance to find a balance between the skills of the animal and the demands of the task when using interactive dog games. The use of this type of game is an opportunity to combine benefits of human-dog interaction, feeding and cognitive enrichment.

## Acknowledgements

First, we thank the staff of the animal shelter for supporting the study and TRIXIE Heimtierbedarf GmbH & Co. KG for providing the interactive game used in the study. Furthermore, we would like to thank Andreas Futschik who provided statistical support, Christian Haberl for his technical support, Jean-Loup Rault for his support in error correction of RR data and Zsófia Virányi for commenting the manuscript with regard to aspects of cognition.

