## Supplemental material for "Balancing skill against difficulty - behavior, heart rate and heart rate variability of shelter dogs during two different introductions of an interactive game"

**Table S1** Types of errors and methods for their identification and correction.

| ***error type*** | **Description** | **identification** | **correction** | **reference** |
|---|---|---|---|---|
| ***Type 1*** | IBI value < 900 ms or value between 900 and 1200 ms and both normal surrounding IBIs, multiplied by two, are bigger than the deviating value | visual and/or IBI difference greater than 450 ms | replace by the mean of the nearest two surrounding correct values | (Marchant-Forde et al. 2004) |
| ***Type 2*** | very small IBI value followed by a very large IBI value | visual and/or IBI difference greater than 450 ms | replace by the mean of the nearest two surrounding correct values | (Gamelin et al. 2006, 2008, Giles et al. 2016) |
| ***Type 3*** | very large IBI value followed by a very small IBI value | visual and/or IBI difference greater than 450 ms | replace by the mean of the nearest two surrounding correct values | (Gamelin et al. 2006, 2008, Giles et al. 2016) |
| ***Type 4(a)*** | IBI value between 900 and 1200 ms and one or both of the normal nearest IBIs, multiplied by two, is smaller than the deviating value | visual and/or IBI difference greater than 450 ms | divide by 2 and insert two new values | (Gamelin et al. 2006, 2008, Giles et al. 2016, Lensen et al. 2017) |
| ***Type 4(b)*** | IBI value between 1201 and 1800 ms and one or both of the normal nearest IBIs, multiplied by two, is smaller than the deviating value | visual and IBI difference greater than 450 ms | divide by 2 and insert two new values | (Gamelin et al. 2006, 2008, Giles et al. 2016, Lensen et al. 2017) |
| ***Type 4(c)*** | IBI value between 1801 and 2700 ms and one or both of the normal nearest IBIs, multiplied | visual and IBI difference greater than 450 ms | divide by 3 and insert three new values | (Gamelin et al. 2006, 2008, Giles et al. |

| *error type* | Description | identification | correction | reference |
|---|---|---|---|---|
| | by three, is smaller than the deviating value | | | 2016, Lensen et al. 2017) |
| ***Type 5*** | several consecutive very low IBI values | visual | sum up the values and replace by the sum | (Gamelin et al. 2006, 2008, Giles et al. 2016) |
| ***consecutive*** | three or more identical consecutive IBI values | IBI difference = 0 | delete all but one value | (Jonckheer-Sheehy et al. 2012) |
| ***delete*** | IBI values > 2700 ms (partially) followed by very small IBI values (e.g. 50 ms) | visual and IBI difference greater than 450 ms | delete the deviating value(s) | (Schöberl et al. 2014) |

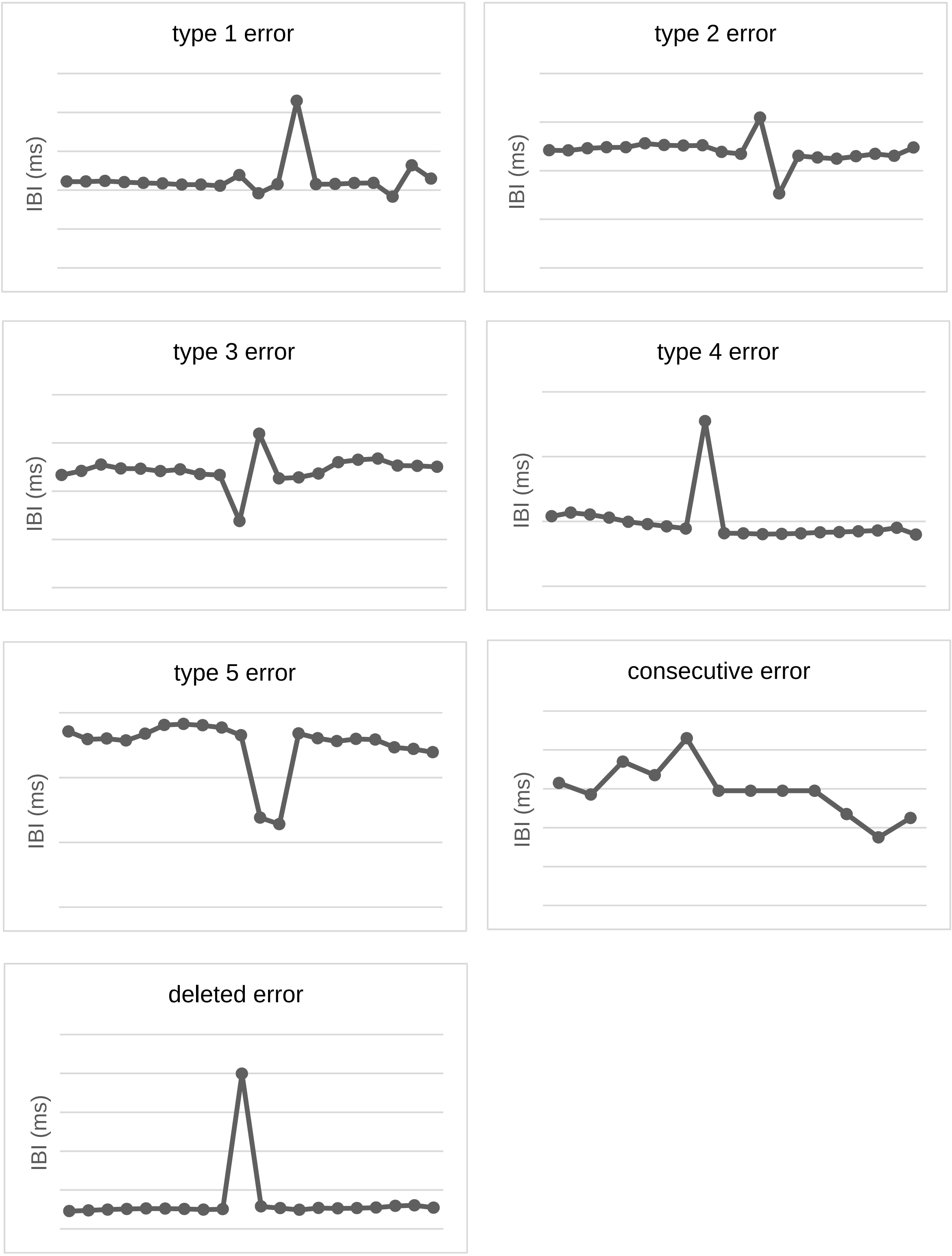


**Fig. S1** Examples for the used error types 1 – 5, consecutive and deleted error from the manual error correction of Polar data in Excel. The figures were created with the data set of this study.

### Decision tree

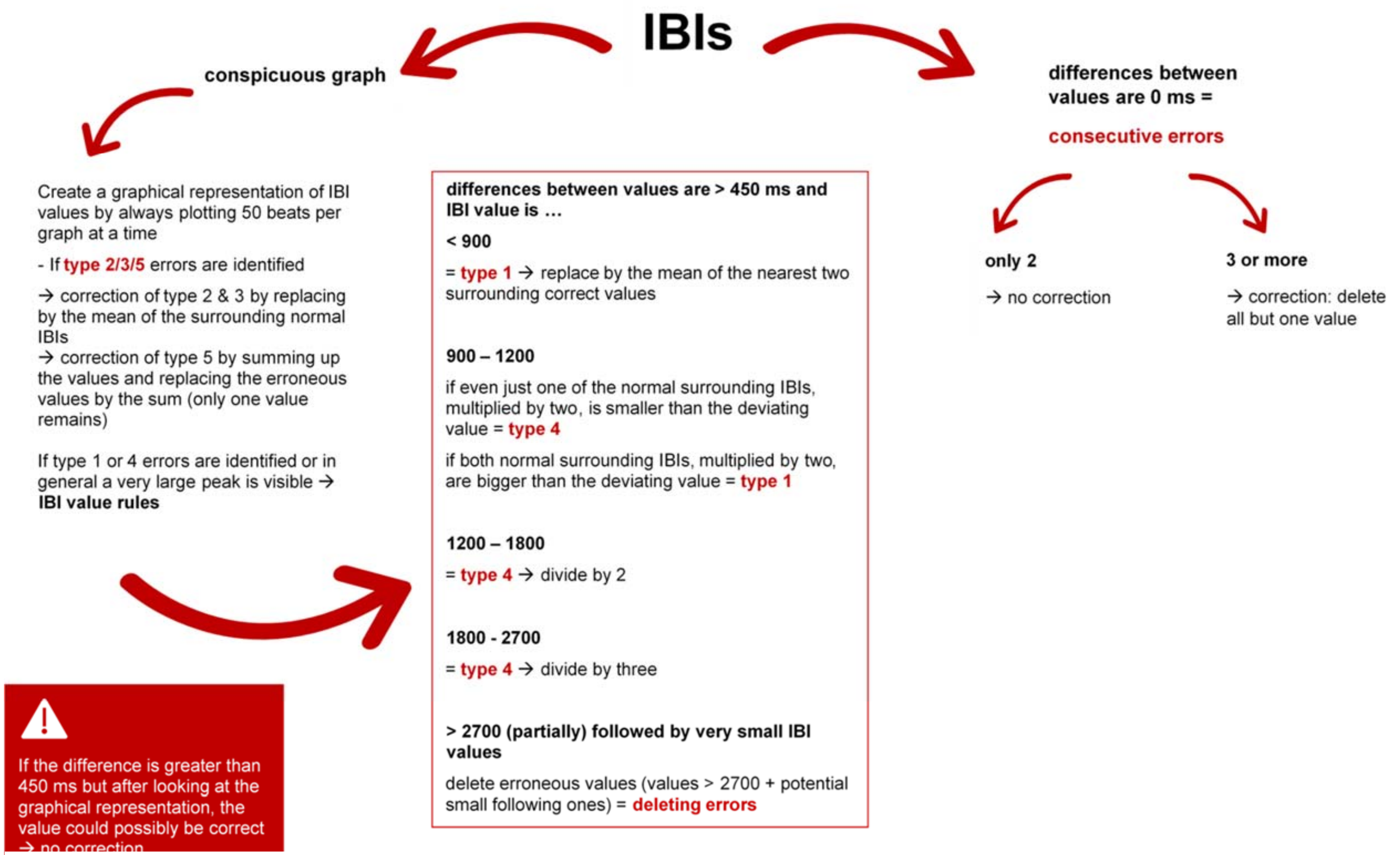


**Fig. S2** Decision Tree for the manual error correction procedure used in this study.
